# Selection constrains high rates of satellite DNA mutation in *Daphnia pulex*

**DOI:** 10.1101/156554

**Authors:** Jullien M. Flynn, Ian Caldas, Melania E. Cristescu, Andrew G. Clark

## Abstract

A long-standing evolutionary puzzle is that all eukaryotic genomes contain large amounts of tandemly-repeated satellite DNA whose composition varies greatly among even closely related species. To elucidate the evolutionary forces governing satellite dynamics, quantification of the rates and patterns of mutations in satellite DNA copy number and tests of its selective neutrality are necessary. Here we used whole-genome sequences of 28 mutation accumulation (MA) lines of *Daphnia pulex* in addition to six isolates from a non-MA population originating from the same progenitor to both estimate mutation rates of abundances of satellite sequences and evaluate the selective regime acting upon them. We found that mutation rates of individual satellite sequence “kmers” were both high and highly variable, ranging from additions/deletions of 0.29 – 105 copies per generation (reflecting changes of 0.12 - 0.80 percent per generation). Our results also provide evidence that new kmer sequences are often formed from existing ones. The non-MA population isolates showed a signal of either purifying or stabilizing selection, with 33 % lower variation in kmer abundance on average than the MA lines, although the level of selective constraint was not evenly distributed across all kmers. The changes between many pairs of kmers were correlated, and the pattern of correlations was significantly different between the MA lines and the non-MA population. Our study demonstrates that kmer sequences can experience extremely rapid evolution in abundance, which can lead to high levels of divergence in genome-wide satellite DNA composition between closely related species.

## INTRODUCTION

Up to half of the genome of higher eukaryotes is composed of tandem arrays of simple repetitive motifs that can span megabases, called satellite DNA (Platero et al. 1998, Padeken et al. 2015). Satellite DNA had initially been thought to carry no useful function, and because it posed a replication burden, it became known as “junk DNA” (Orgel and Crick 1980). It also has the potential to be harmful because it can cause deleterious genomic rearrangements facilitated by recombination between similar motifs on different chromosomes (Bzymek and Lovett 2001). Thus, evolutionary biologists have long wondered why repeated sequences accumulate and are maintained at such abundance in the genome. Satellite DNA is typically found in heterochromatic regions where expression is silenced and recombination is low, in Y chromosomes, and in centromeric and telomeric regions (Charlesworth and Charlesworth 2000; Henikoff et al. 2001). One hypothesis is that being situated in heterochromatin, where recombination is suppressed, could both minimize the cost of repeated DNA and reduce the efficacy of selection against it, allowing its accumulation (Charlesworth et al. 1994). Moreover, the observation that, genome-wide, satellite repeats undergo rapid turnover, with nearly complete replacement of these repeat sequences even between closely related species poses challenging questions about the evolutionary forces that shape their evolution (Subirana et al. 2015; Wei et al. 2014). The study of satellite DNA evolution on the timescales of diverged populations and species have revealed interesting patterns, but understanding the forces that generated these patterns requires direct observation on a shorter timescale.

Early evolutionary models did not consider selection to be influencing satellite sequence evolution, except perhaps selection on overall genome size (MacGregor and Sessions 1986; Charlesworth et al. 1994; Stephan and Cho 1994). Stephan and Cho (1994) proposed a model where tandemly-repeated sequences could be generated and expand from mutation, recombination, and replication slippage, combined with selection to maintain the genome size. Another model, the “library hypothesis,” attempts to explain the differences in satellite composition by posing that the common ancestor of related species contained a library of many satellites, which are then differentially amplified within each lineage as they diverge (Fry and Salser 1977). However, the rate at which satellite sequences expand and contract is difficult to quantify. Most models pose evolutionary neutrality of satellite repeats, but we now know that satellite sequences also carry vital cellular functions. Since much of satellite DNA is located in centromeric regions, there is evidence that selection acts on the sequence motifs themselves, as they serve to bind centromere and histone proteins for centromere maintenance and chromosome separation (Henikoff et al. 2001; Malik 2009). These functions have implications for genome stability, and changes in satellite DNA sequences have been shown in some cases to be drivers of speciation by inducing chromosome rearrangements leading to karyotype divergence (Paco et al. 2015). Moreover, some satellite sequences are transcribed and may be involved in the regulation of heterochromatin formation (Palomeque and Lorite 2008; Plohl et al. 2008). Even more intriguing is the observation that perturbation of satellite content can alter gene expression genome-wide (Lemos et al. 2008). At present, it is not known whether these examples of functional importance of satellite DNA are the exceptions or the rule and to what extent selection governs the evolution of satellite DNA.

Here, we refer to the units or words of tandemly repeated satellite sequences as “kmers”. Several potential and non-mutually exclusive selection regimes could be operating on kmer arrays: negligible selection where satellite DNA content is primarily governed by mutation and drift, stabilizing selection to maintain a particular “optimal” composition, negative selection to purge satellite DNA, and positive selection for rapidly generating new kmer motifs. Studying satellite DNA changes over an evolutionary timescale showed clear differences across geographic subpopulations of a single species and almost complete turnover between species, confirming that they evolve rapidly (Wei et al. 2014; Subirana et al. 2015). However, quantifying the relative contributions of mutation, genetic drift, and selection is difficult since all these forces are at play in influencing genetic variation in natural populations. Additionally, it has been a challenge to quantify the genome-wide satellite composition since their repetitive nature makes them problematic to sequence and assemble (Hoskins et al. 2007). Early studies describing satellite composition in different species have focused on single satellites or a small family of satellite sequences (Lohe and Brutlag 1987). Obtaining a genome-wide view of the rate of mutation in satellite sequences and how selection shapes their evolution would facilitate the understanding the longstanding puzzles of satellite DNA evolution.

Mutation accumulation (MA) experiments reduce selection to a minimum by enforcing bottlenecks every generation and reducing the effective populations size so that both neutral and deleterious mutations can accumulate almost neutrally (Simmons and Crowe 1977). An MA experiment combined with observations from a population with the same genetic background where selection is not removed allows the study of the effects of mutation alone versus the effects of mutation combined with selection (Flynn et al. 2017). *Daphnia pulex* is an ideal organism in which to study satellite DNA mutation, firstly because MA studies can be conducted effectively under asexual reproduction and single-progeny bottlenecks. Short microsatellite arrays have been studied in *D. pulex* (Seyfert et al. 2008; Sung et al. 2010), but the sequence motifs and abundances of satellite DNA have not been characterized, although 25% of the genome is estimated to be heterochromatic (Colbourne et al. 2011). *D. pulex* has a potentially dynamic genome, with evidence of high rates of deletion and duplication (Keith et al. 2016). The lineage we use reproduces exclusively asexually via apomixis, meaning changes in satellite content represent spontaneous mutations in the germline without the opportunity for meiotic drive to play a role (Malik 2009; Wei et al. 2017).

In this study, we compare the kmer composition of MA lines (without selection) to individuals raised in a competitive non-MA population (with selection) of the identical *D. pulex* genetic background. We were interested in exploring the mutational dynamics of satellite DNA that can often not be reliably mapped to the reference genome. We quantify the kmer composition using a mapping-independent method to capture all tandem kmers up to 20 bp detectable from short-read data, without introducing biases from the reduced representation of repetitive sequences in the reference genome. Our approach allows us to (1) estimate the kmer mutation rates, including expansions, contractions, complete loss of kmers, and generation of new kmers; and (2) evaluate the type of selection (if any) acting on the satellite DNA sequences. Given the observed natural variation in satellite DNA content among populations and species, we expect expansions and contractions to occur at high rates in the MA lines. If satellite DNA is under selective constraint, we expect there to be less variation in kmer abundance in the population evolving under selection than the neutrally-evolving MA lines.

## MATERIALS AND METHODS

### Daphnia line setup and DNA sequencing

The current study includes a total of 34 genomes: 24 MA line genomes and 6 population isolate genomes that were sequenced in Flynn et al. (2017), and four additional previously unpublished MA genomes. The asexually reproducing MA lines and a non-MA population were initiated from a single progenitor (see Flynn et al. 2017 for a detailed description of the MA experiment). MA lines were propagated for an average of 82 generations before whole-genome sequencing with single-individual bottlenecks between generations prior to DNA isolation and whole-genome sequencing. In contrast, the non-MA population was maintained without inducing bottlenecks for 46 months in a 15 L tank, before six individuals were isolated for sequencing. The approximate census population size was estimated to be 100-250. Overlapping generations occurred so the exact number of generations that the population isolates progressed could not be recorded. Thus, we used a life history experiment and estimated the slowest and fastest moving lineages to calculate that the population underwent at least 62 generations (Supplementary Material File S2). Prior to sequencing, *Daphnia* individuals were subjected to a brief antibiotic treatment and fed with sterile Sephadex beads to reduce contaminants before sequencing (Fields et al. 2015).

Sequencing libraries were prepared with Illumina Nextera procedures in two batches and all genomes were sequenced to approximately the same coverage (10x). To ensure reproducibility between multiple library preparations, we performed technical replication on two MA lines, such that two independent library preparations and Illumina sequencing runs were done for the MA lines C01 and C35 (in a completely separate, third sequencing batch). We analyzed independently all technical replicates to ensure that variation produced by different library preparations is smaller than the variation due to biological expansion/contraction mutations. Libraries were sequenced on an Illumina HiSeq 2000 instrument at Genome Quebec of McGill University with 100 bp paired-end reads.

### Satellite quantification

To remove redundant sequences, adapters were trimmed and overlapping reads were merged with SeqPrep (https://github.com/jstjohn/SeqPrep). To identify and quantify satellites, the resulting unmapped read files were used as input for the program k-Seek (Wei et al. 2014, https://github.com/weikevinhc/k-seek). The current version of k-Seek detects words or “kmers” of length 1-20 bp that are repeated tandemly to cover at least 50 bp within the same read. The program allows for one single nucleotide mismatch per repetition of each kmer (see Wei et al. 2014 for details). All offsets and reverse complements of each kmer are compiled and the total of all individual unit copies are summed across all reads. Kmer sequences are presented as the strand/offset that is alphabetically ordered (i.e. A’s will be at the start of the sequences if possible). This method of tandem kmer detection and quantification has been found to be reproducible across different library preparations (Wei et al. 2014).

To obtain a quantitative comparison between samples we normalized the kmer counts by both individual library sequencing depth and GC content, as PCR-based library preparations are known to show a bias in the GC content of the fragments amplified and sequenced (Benjamini and Speed 2012). This is especially problematic for satellite analysis since, by nature, many of the repeated sequences may have extreme GC contents. First, reads were mapped against the *Daphnia pulex* reference genome (version 1, Colbourne et al. 2007) using BWA (Li and Durbin 2009) v0.7.10 with default settings. Output BAM files spanning the whole genome were given as input to a custom shell script (https://github.com/jmf422/Daphnia-MA-lines/tree/master/GC_correction) to calculate correction factors following Benjamini and Speed (2012). The correction factors produced for high and low GC contents were extreme and variable, which was likely due to low read counts giving unstable estimates as well as mapping biases to the reference genome (Flynn et al. 2017). Thus, to smooth the correction, we used a Python script to bin values of GC content together, employing wider bins for GC content < 0.25 and > 0.60 (https://github.com/jmf422/Daphnia-MA-lines/tree/master/GC_correction). The correction factor was then applied to kmer counts according to which GC bin their content belonged.

### Mutation and interspersion metrics

We define mutations in kmer repeats as any change in the number of copies of each specific kmer. The rate of mutation is then the observed change in the number of each kmer per generation. Each kmer may be in tandem arrays found in one or in several locations across the genome, and our method sums the counts of each kmer across all genomic regions. A mutation rate for a given kmer reflects the sum of changes in copy number across these potential multiple loci. To calculate the mutation rates, we used the mean abundance of the kmer in the non-MA population isolates as a proxy for the ancestral abundance using the following equation:

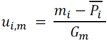

Where *u*_*i, m*_ is the mutation rate of kmer *i* in MA line *m, m_i_* is the abundance of kmer *i* in MA line 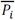 is the mean abundance in the population for kmer *i*, and *G*_*m*_ is the number of generations propagated for MA line *m*. *u* could be negative (for contractions) or positive (for expansions). We used the same equation to calculate the absolute rate of change of each kmer, except we took the absolute value of the numerator.

As mutation rates could be correlated with copy number, we also calculated the absolute mutation rate normalized by initial copy number using the following equation:

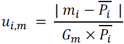

We used the same equations for calculating the “realized” kmer mutation rates (mutations that made it through selection in the population isolates), and we used the conservative estimate of 62 for the number of generations.

We also searched for new kmers generated *de novo* in the MA experiment. In order to be considered as a new kmer, putative new kmers had to have at least 3 copies in at least one MA line and be completely absent from all other lines. New shared kmers had to have at least 3 copies in one MA line and at least 2 copies in a second line. Since k-Seek has a detection limit for kmer arrays that are at least 50 bp long, we checked if the putative new kmers were present in shorter arrays (~25 bp) in the other MA lines to determine if the kmer sequence was generated truly de novo or if it expanded from an already present repeated motif. Similarly, we searched for kmers that had been lost in individual and pairs of MA lines throughout the course of the experiment. Our criteria for identifying lost kmers were that they had to have 0 copies in the affected line or pair of lines, and at least 2 copies in all other lines. We checked if putative lost kmers were completely undetectable or if they were still present in short arrays (~25 bp), below the detection threshold of k-Seek.

Since we have paired-end reads, we have the potential to detect the level to which kmers are present on both reads of the pair (on the same genomic fragment). To quantify this interspersion level, we used the metric *I*, as in Wei et al (2014):

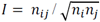

where *n*_*ij*_ is the number of read pairs containing kmer *i* and kmer *j*, and *n*_*i*_ and *n*_*j*_ are the number of read pairs containing kmer *i* and kmer *j* respectively.

### Data availability

Raw sequence files have been deposited in the NCBI Sequence Read Archive BioProject ID: PRJNA341529 BioSamples SAMN05725816 and SAMN05725817. All analyses were performed using RStudio (V 0.99.903), Python V2 and Perl V5 scripts. Html files showing the code and output to all the analyses are available in the Supplementary Material, and all scripts as well as the required processed input files are available at https://github.com/jmf422/Daphnia-MA-lines. File S1 contains descriptions of all supplementary files, as well as the supplementary table and figure legends.

## RESULTS

### Library and GC correction

Over 713 million unmapped reads across all samples were scanned for tandemly-repeated sequences with k-Seek. Of these, over 5.85 million reads (0.82 %) were found to be composed of kmers of unit length 1-20 bp and encompassed at least 50 bp of the read. Although the second library batch had overall more reads, a similar proportion of the total reads contained kmers in both libraries (Supplementary Figure S1). Unless otherwise noted, the abundance of each kmer is presented in copies per 1x read depth after GC normalization.

We found that our qualitative conclusions were robust whether or not we applied a correction factor to normalize the kmer counts based on each kmer’s GC content. Our results were also robust across a wide range of parameters used for the GC correction. As library preparations involving PCR have been shown to result in a biased representation of fragments based on GC content (Benjamini and Speed 2012), we present results after GC correction as described in the Materials and Methods.

Since the libraries were prepared in two separate batches, we were concerned that this could confound our results. A Principal Components Analysis (PCA) was able to separate samples based on library batch along PC2, but PC1 appeared to separate samples based on biological mutation patterns (Supplementary Figure S2). There were no consistent differences across all kmers or GC contents of kmers that could be consistently corrected for (Supplementary Material File S4). In order to ensure that library batch would not be a confounding factor, we performed a technical replicate of two MA lines and showed that the abundances of each kmer between the library prep batches were highly similar (Supplementary Material File S4). Although there was minor variation between technical replicates in some kmers, their overall mutational patterns were highly similar, shown by their high clustering on PC1 (Figure S2). Additionally, we tested for differences between the mean abundances of the kmers between the MA lines prepared in the first batch and the MA lines prepared in the second batch. The mean abundances of most of the kmers (28 out of 39) in the second-batch MA lines were within the 99% confidence intervals from 1000 subsampling replicates of four first-batch MA lines. 30 of the kmers had mean abundances in the second batch within the range of observed abundances of the first-batch MA lines (Supplementary Material File S4). For these reasons, we did not consider library preparation batch to be a significant confounding factor.

### Description of satellite DNA content in *Daphnia pulex*

We first sought to describe and quantify the genome-wide short repeat content in *Daphnia pulex* and make inferences about kmer origins. There were 162 kmers that had at least 2 copies per 1x coverage in all our *D. pulex* lines. There were 39 kmers that had an average abundance of at least 100 copies, and 12 that had an average abundance of at least 1000 copies after normalization (Table S1). We chose to focus most of our analysis on the 39 kmers that had at least 100 copies after normalization (Figure 1). The most abundant kmer, the poly-A repeat, was present at an average of 79,528 copies in each genome. The second most abundant kmer, present at an average of 62,281 copies, was AACCT. This 5-mer is known to be the ancestral telomere repeat in Arthropods (Sahara et al. 2009) and was previously found in *Daphnia* (Colbourne et al. 2011). Runs of poly-C were also found to be in the “top 39 kmers”, having an average of 6379 copies. Two of the four possible 2-mers were included, AG and AC, having an average of 408 and 244 copies, respectively. The most abundant kmer sizes were 5-mers (there were six 5-mers of the most abundant 39 kmers), 10-mers (7/39), and 20-mers (7/39). No 15-mers were abundant (>100 copies) in the genome, but there were 13 among all 162 kmers. Overall, 20-mers were the most represented with 50 kmers of the 162. Adding up the total short kmer content per genome in base pairs, we found the median to be 1.20 Mb per 1x coverage, which represents 0.6 % of the estimated 200 Mb genome.

**Figure 1:**
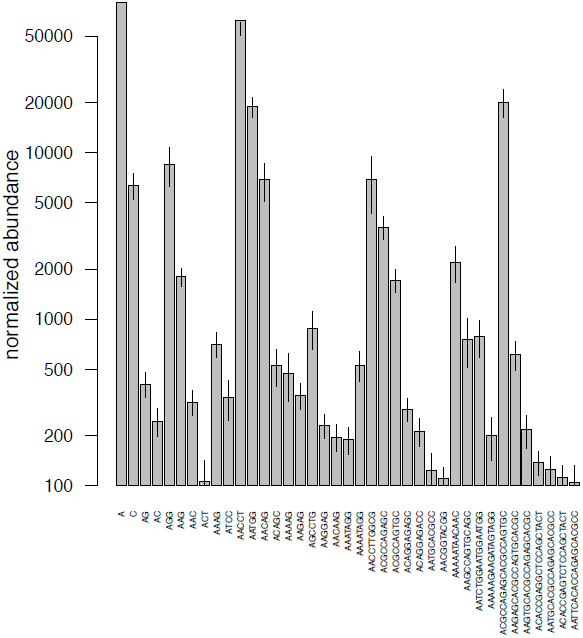
Abundance of the kmers that contain at least 100 counts after normalization. Kmers are arranged in order of length, from 1-mers to 20-mers.

We also compared the satellite composition in *D. pulex* to that in *Drosophila melanogaster*, the only other arthropod that has had its genome-wide satellite content characterized to the same level of detail (Wei et al. 2014). This comparison might indicate the extent of satellite diversity across arthropods and functional conservation of some satellites. Comparing the most abundant kmers (at least average 100 copies normalized) between these two species, we found that 10 short kmers (mostly derivations of AG repeats) were present in both species: A, C, AC, AG, AAC, AAG, ACT, AAAAG, AAGAG, and AAGGAG. We found several short (3-5 bp) motifs to be highly represented in the most abundant kmers, and among the 162 kmers (Table 1). Most of these motifs were completely absent or rare within the top 108 *D. melanogaster* kmer sequences, except for AG-type motifs (Table 1).

**Table 1.** Number of kmers that contain certain sequence motifs. The top 39 kmers are characterized by having an average of at least 100 copies per 1x coverage, and the top 162 kmers have at least 2 copies per 1x coverage in all samples. The *D. melanogaster* kmers are the 108 kmers that have at least 100 copies normalized (personal communication, K. Wei).

| Motif sequence | In top 39 kmers | In top 162 kmers | In *D. melanogaster* kmers |
| --- | --- | --- | --- |
| AAAA | 4 | 13 | 12 |
| AAC | 7 | 32 | 16 |
| AAG | 11 | 49 | 45 |
| ACGC | 8 | 52 | 0 |
| AGC | 12 | 69 | 4 |
| AGG | 11 | 41 | 14 |
| AGGAG | 3 | 15 | 5 |
| GCCAG | 7 | 48 | 0 |
| TAGG | 4 | 10 | 0 |
| TCCAG | 3 | 14 | 0 |

In order to understand the origins and dynamics of kmer sequences in *Daphnia*, we inspected the kmers for sequence similarities. We found that many kmers belonged to families of related sequences separated by a small number of potential mutational steps. Figure 2 shows a network diagram highlighting the potential mutational relationships between 29 of the 39 of the top kmers. The most striking example of a family of related kmers is the 9 kmers of length 10-20 bp shown in the top left corner of Figure 2. Ten kmers were not included in the network, either because they were mono-, di-, or tri- nucleotides whose sequence similarities were not meaningful (6 kmers: A, C, AC, AG, AGG, ACT), or because they did not have a clear mutational relationship with any of the other top kmers (4 kmers: AGCCTG, ACAGC, ATCC, AACGGTACGG).

**Figure 2:**
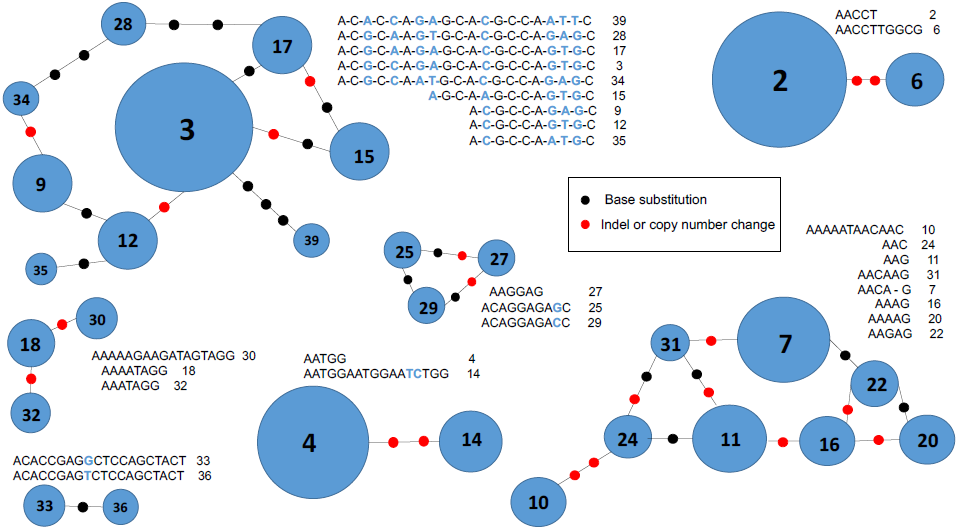
Network of relatedness of the 29 of the 39 most abundant kmers. Networks show possible mutational steps between related kmers. Kmers are numbered based on their abundance rank. The sizes of the circles also correspond to the kmer abundance.

### Extremely high rate of mutation in satellite repeats

Because we could not call single nucleotide changes reliably without mapping to the reference genome, we were not able to estimate mutation rates for single base changes in the kmers, and restrict our attention to changes in copy number. To estimate the copy number mutation rates, we used the mean abundance of each kmer in the population isolates as a proxy for the ancestral abundance of the kmer in the MA progenitor. This should be a reasonable estimation assuming that individuals in the population roughly maintained the ancestral kmer content (see below). In fact, all kmers were similar in abundance between the MA lines and the population isolates: all kmer abundances in the population were within 2 standard deviations of the abundance in the MA lines, and 69% were within 1 standard deviation. This is consistent with the MA lines both gaining and losing kmer copies. The absolute mutation rates (summing expansions and contractions) per kmer sequence ranged from 0.29 to 105 copies/generation, with a median of 1.26 copies/generation. This is not including the telomere repeat, which had the highest rate of change at 199 copies/generation. It is likely that the changes affecting the telomere repeats are from a different, non-mutational process (i.e. caused by a mechanism involving telomerase which is not transgenerationally inherited). The suppression of meiosis in these *Daphnia* lines may have resulted in relaxed selection on telomere maintenance, but we have no direct evidence in support of this.

We found that the mutation rates were correlated with the kmer’s initial copy number abundance, proxied by the mean abundance of the population isolates, and that the best fit of this relationship is linear (*r*^*2*^ #x003D; 0.88, *p* < 2.2 ×10^-16^). Thus, to compare mutation rates of individual kmers, we normalized the mutation rates by the initial unit copy number to give mutation rates in copies per generation per original copy. Even after this normalization, we still found mutation rates to be exceptionally high – on the order of 10^-3^ per copy per generation – compared to previous estimates of other types of mutation such as microsatellite mutations. Normalized mutation rates ranged from 1.23 × 10^-3^ to 7.97 × 10^-3^, with a mean of 2.74 × 10^-3^ copies per generation per original copy. Expansions were more frequent than contractions, with more MA lines experiencing expansions in most kmers, and the overall rate of change was in a positive direction for 124/162 kmers (76%) (Figure 3a and 3b). Expansion and contraction rates were not correlated with either GC content or kmer size (Supplementary Material Figure S3).

**Figure 3:**
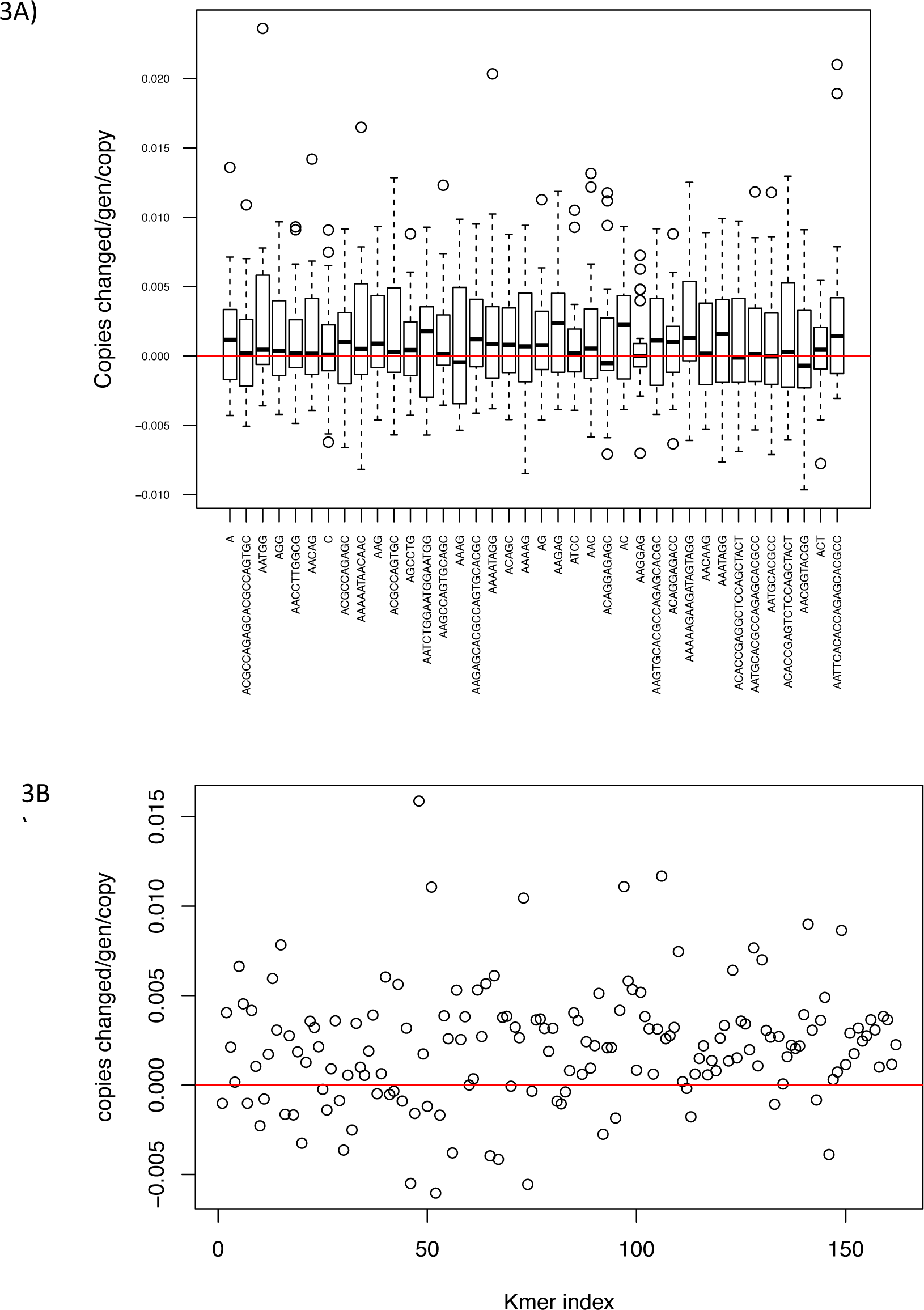
Kmer mutation rates. (A) Boxplot of mutation rates in the MA lines (n=28). The top 20 kmers are shown, in descending order of abundance. Negative values indicate contractions and positive values indicate expansions. (B) Plot of mean overall mutation rate across all 162 kmers at normalized abundance of at least 2 copies per 1x coverage (in descending order of abundance). The red line indicates an overall mutation rate of 0.

A multi-generational MA experiment provides the potential opportunity to observe completely new kmer sequences arising. We searched for kmers that were generated *de novo* during the experiment, in single lines and shared across pairs of MA lines. We found five kmers gained in single MA lines and three gained in three independent pairs of MA lines (Table 2). These MA lines did not share single nucleotide mutations within uniquely mapping regions of the genome (Flynn et al. 2017). Five of the eight new kmers seemed to originate from a single base substitution of an existing kmer present in high abundance (Table 2). Two of the other new kmer sequences contained motifs that were abundant in existing kmers. Only 3 out of 8 putative new kmers were present in shorter arrays (~25 bp, half of k-Seek’s requirement) in the ancestral lineage (Table 2). We also searched for kmers that were lost from individual and pairs of MA lines. There were seven kmers lost uniquely and two lost in a pair of lines (Table 3). Three of the seven lost kmers were lost from MA line C40. C40 is the MA line that was found to have ~100 kb of homozygous deletions of non-repetitive sequence mapped to the reference genome on one chromosome (Flynn et al. 2017), so it is possible that these lost kmers were present exclusively in the deleted regions. Of the lost kmers, 5 out of 7 were still detectable in shorter arrays (~25 bp) in the affected MA line (Table 3).

**Table 2.**
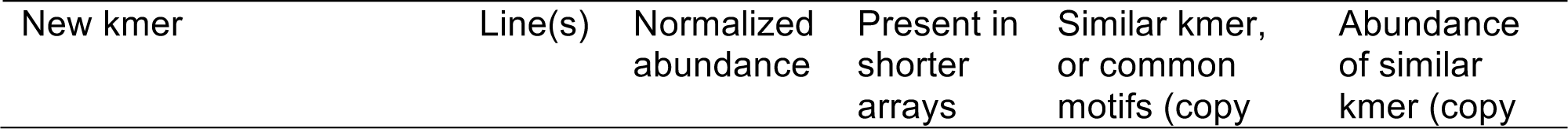
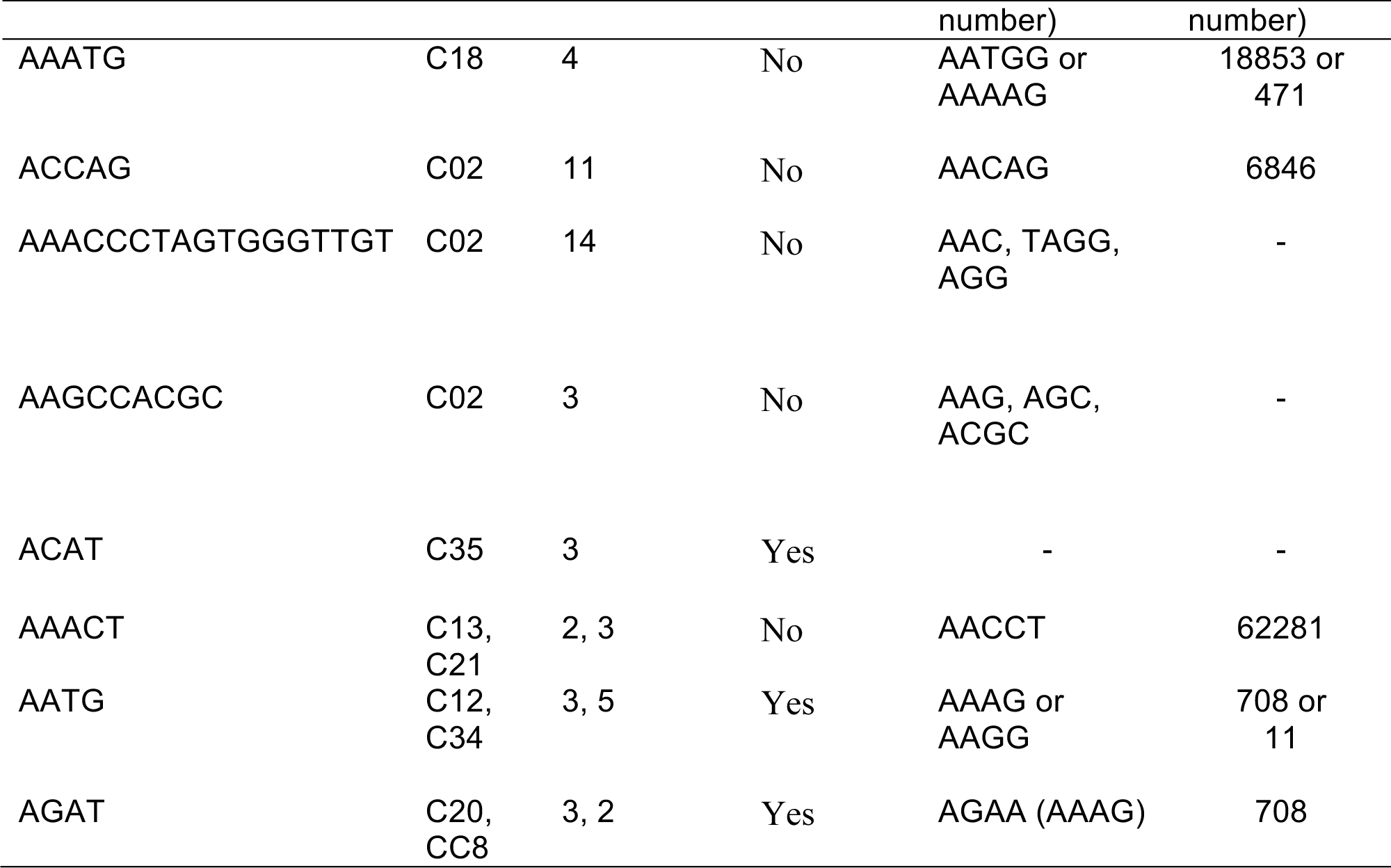
New kmers generated *de novo* during the MA experiment. Many of them are potentially generated from single nucleotide mutations or rearrangements of existing kmers. It is indicated if the putative new kmer was present in shorter arrays (~25 bp) in the ancestral lineage.

**Table 3.**
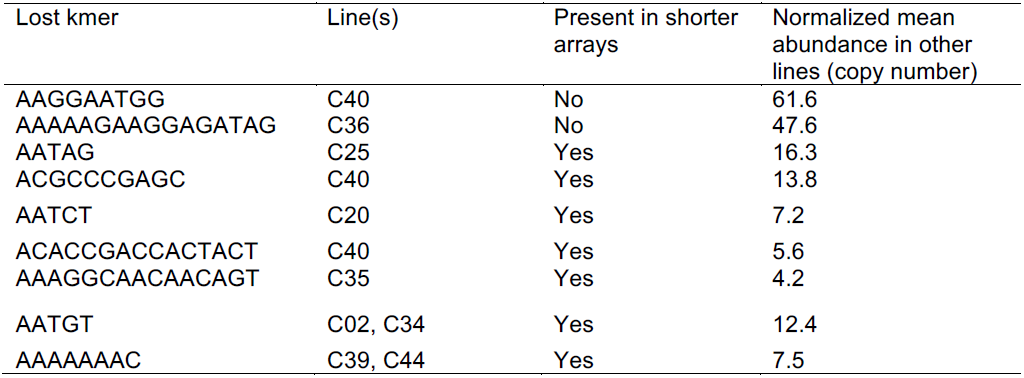
Kmers that were lost during the course of the MA experiment. It is indicated if the putatively lost kmer is present in short arrays at low frequencies in the MA line in question.

### MA lines versus population with selection

Comparing the changes in kmer composition between the MA lines and the population isolates, which experienced selection, reveals the potential influences of selection on satellite DNA. The population provides a valid comparison to the MA lines because it was previously shown to show signs of purifying selection in both single nucleotide mutations (Flynn et al. 2017) and copy number variants (CNVs) (Chain et al. 2017, in prep). The population did not experience a recent bottleneck and in fact the most recent common ancestor of the isolates analyzed here was the progenitor of the experiment (Flynn et al. 2017). We also made a conservative estimate of the number of generations the population underwent in order to compare mutation rates (Supplementary File S2). First, we compared the total genome-wide amount of tandem repeats by multiplying the length of the kmer with its number of copies in each genome. The MA lines diverged by a factor of 1.67 in overall kmer abundance, from 1.00 Mb to 1.67 Mb total, with a median of 1.23 Mb. The population isolates had both a lower genomic amount of kmers and a narrower range among isolates, ranging from 0.97 to 1.22 Mb, with a median of 1.05 Mb (Figure 4a). A Levene’s test did not detect a significant difference in the variances between these two groups (p #x003D; 0.12). However, the mean total abundance (but not the variance) of kmers in the population was below the 5% confidence interval produced from 1000 replicates of 6 randomly-sampled MA lines (Supplementary Material File S3).

**Figure 4:**
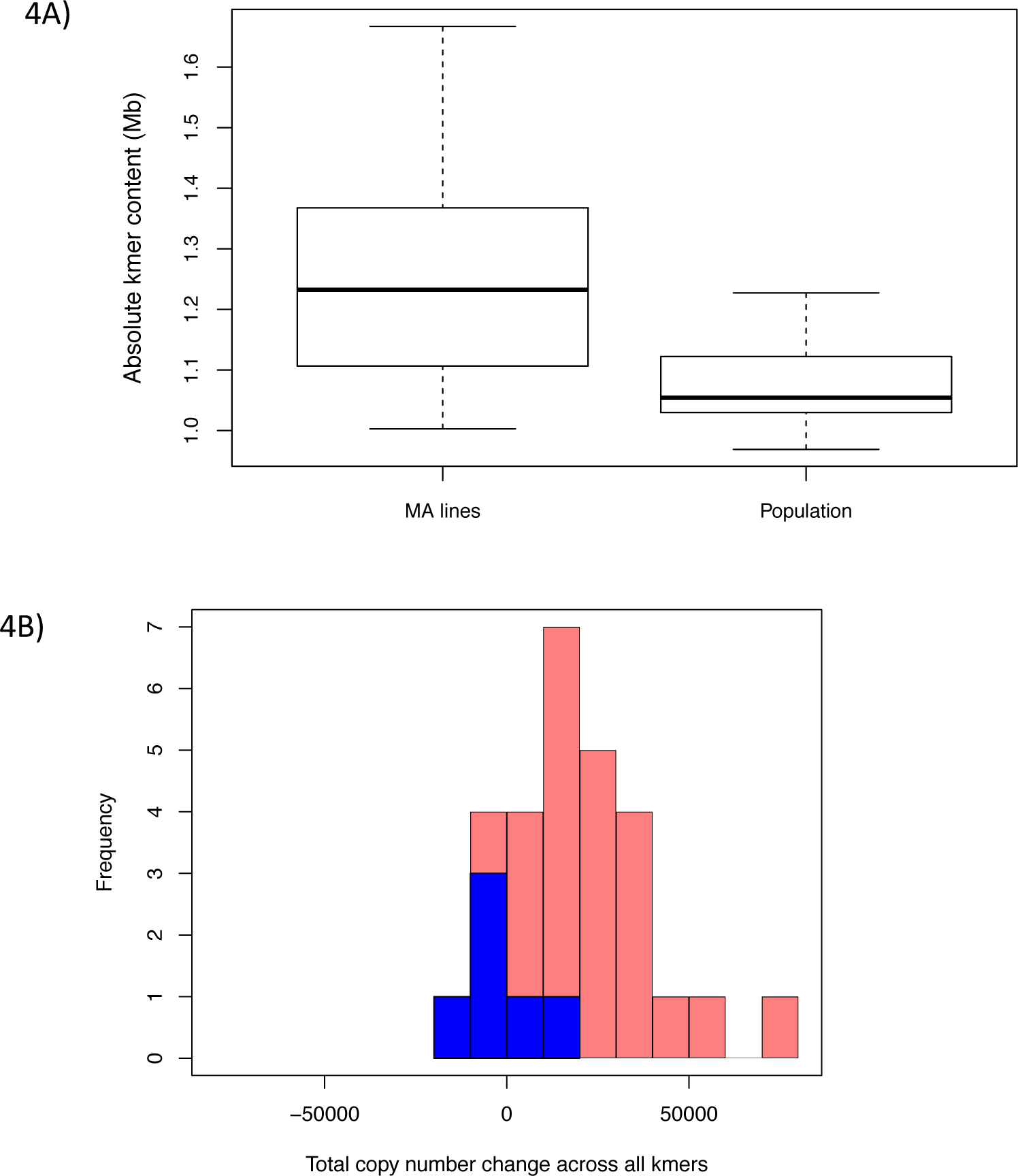
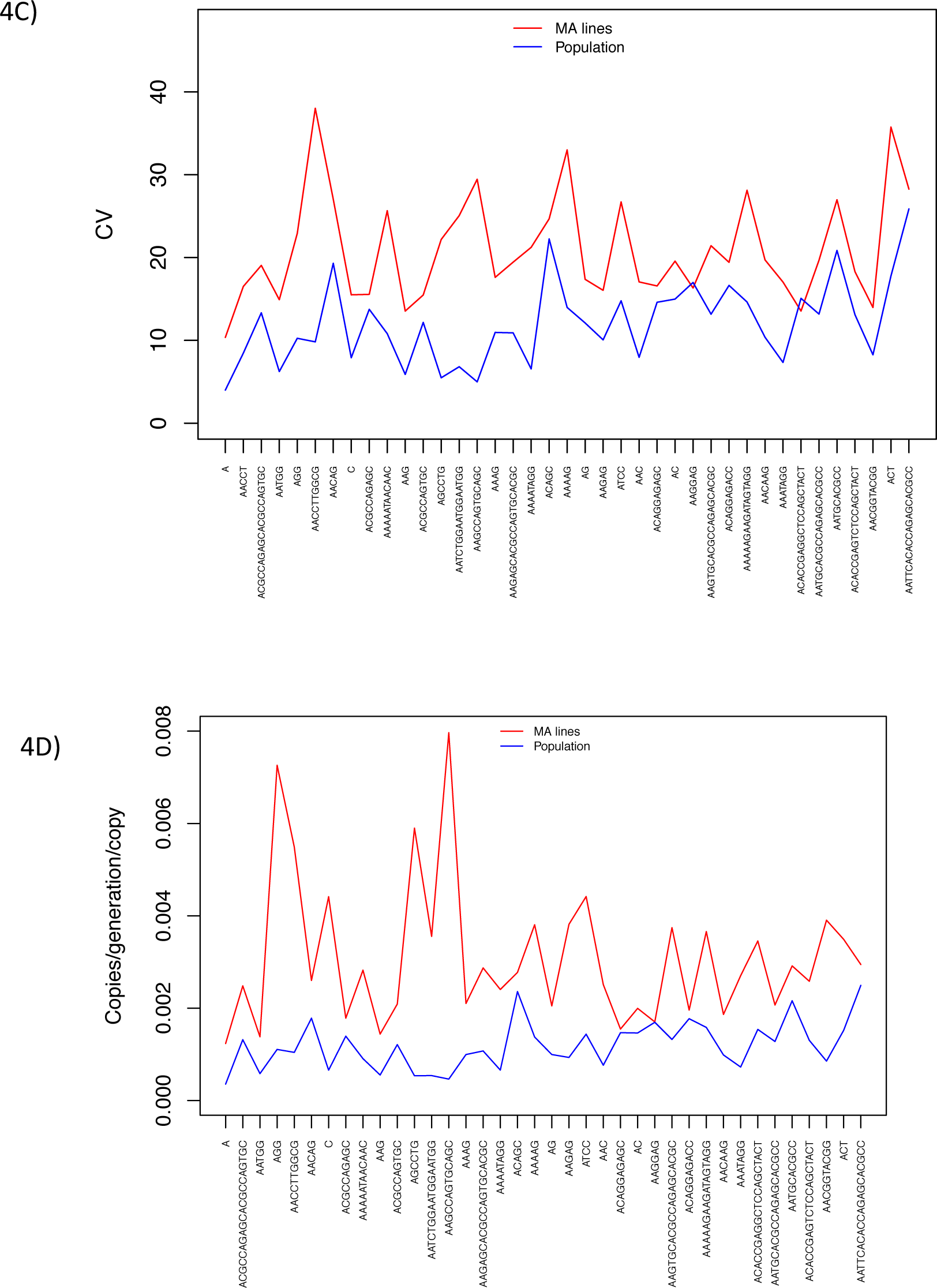
Reduced variation in kmer content in the population compared to the MA lines. (A) Total absolute amount of tandemly-repeated satellite sequences with length 1-20 bp in the MA lines versus the population, including all 162 kmers with at least 2 copies per 1x coverage per line. (B) Histogram of the cumulative copy changes across the top 39 kmers in the MA lines and population. (C) The coefficient of variation across the top 39 kmers for the MA lines and the population. Kmers are ordered by abundance. (D) The absolute normalized mutation rate (sum of expansions and contractions), divided by the initial copy number of the kmer. Data from the MA lines are shown in red and the population in blue.

The MA lines underwent expansions across many kmers, with some MA lines experiencing a considerable increase in satellite DNA content in their genome over the course of the MA experiment compared to the non-MA population isolates (Figure 4b). Most notably was MA line C20, which experienced expansions in 34 of the 39 top kmers, with an overall increase in 580.2 kb in satellite content over 81 generations. On the other hand, population isolates deviated less from an overall balance of expansions and contractions (Figure 4b).

To perform a contrast in the variation of individual kmers between the MA lines and the population, we calculated the coefficient of variation (CV, the standard deviation divided by the mean) for each kmer for both the MA lines and the population. We found that the CV was higher in the MA lines than the population for 37 of the top 39 kmers (Figure 4c, *P* < 2.84 ×10^-9^, sign test). We considered the possibility that the CV of the MA lines was artificially inflated if the mean abundances of some kmers in the second library preparation MA lines were different that of the first library preparation MA lines. We also considered that the CV may be greater in the MA lines partially because the MA lines may have undergone more generations than the population. In order to ensure these factors were not founding our results, we also calculated the CV of the MA lines comparing only the ones that were prepared in the same library and normalized the CV of the MA lines by the difference in the generation number between the MA lines and the population. After this conservative calculation, the CV was higher than the population isolates across 25 of the 39 top kmers (Supplementary Material File S3). Next, we compared the realized mutation rate – the mutations that made it through the “filter” of selection in the non-MA population – to the mutation rate in the MA lines. We used the conservative calculation of 62 generations propagated in the population (Supplementary Material File S2). The realized mutation rates in the population were lower than the MA lines for 32 of the top 38 kmers not including the telomere repeat (Figure 4d, *P* < 2.43 × 10^-5^, sign test). Since the sample size of the non-MA population isolates is lower than the sample size of the MA lines (6 versus 28), we performed 1000 subsampling replicates for each kmer by sampling 6 random MA lines and calculating the mean mutation rate. We found that 22 out of 38 kmers had a lower realized mutation rate in the population than the 5% quantile of the 1000 MA line subsample replicates (Supplementary Material File S3). Moreover, some kmer sequences seemed to be more constrained than others. We roughly quantified constraint by the difference between the mutation rate of the MA lines to the realized mutation rate of the population for each kmer. Five kmers experienced high levels of constraint by this measure, with the realized mutation rate being at least 75% lower in the population than the MA lines: AGG, C, AGCCTG, AAGCCAGTGCAGC, and AATCTGGAATGGAATGG. All of these highly constrained kmers had statistically significant lower realized mutation rates by the subsampling test above. On the other hand, 9 kmers seem to have experienced little constraint in their expansions and contractions, with their realized mutation rates in the population being either slightly higher in the population or very close to that of the MA lines (Figure 4d, Supplementary Material File S3).

Wei et al. (2014) noted strong patterns of correlation of kmer abundances across different populations of *Drosophila melanogaster*, suggesting some sort of evolutionary non-independence. These correlation patterns may suggest constraints on the mutational process, or they may be driven by selection. To investigate correlations in the expansions/contractions between kmers, we computed a correlation matrix (using the Pearson correlation) between each kmer for the MA lines and the population. This would indicate correlations between changes in kmer content inherent to mutation as well as correlations caused by similar selective regimes among kmers. We found that some (mainly positive) correlations existed in the MA lines, but the correlations between kmer changes in the population were more common and stronger for both positive and negative correlations (Figure 5a and 5b). A Mantel test indicated that these correlation matrices were significantly different (*P* < 0.001), indicating a role of selection in interactions between satellite sequences.

**Figure 5:**
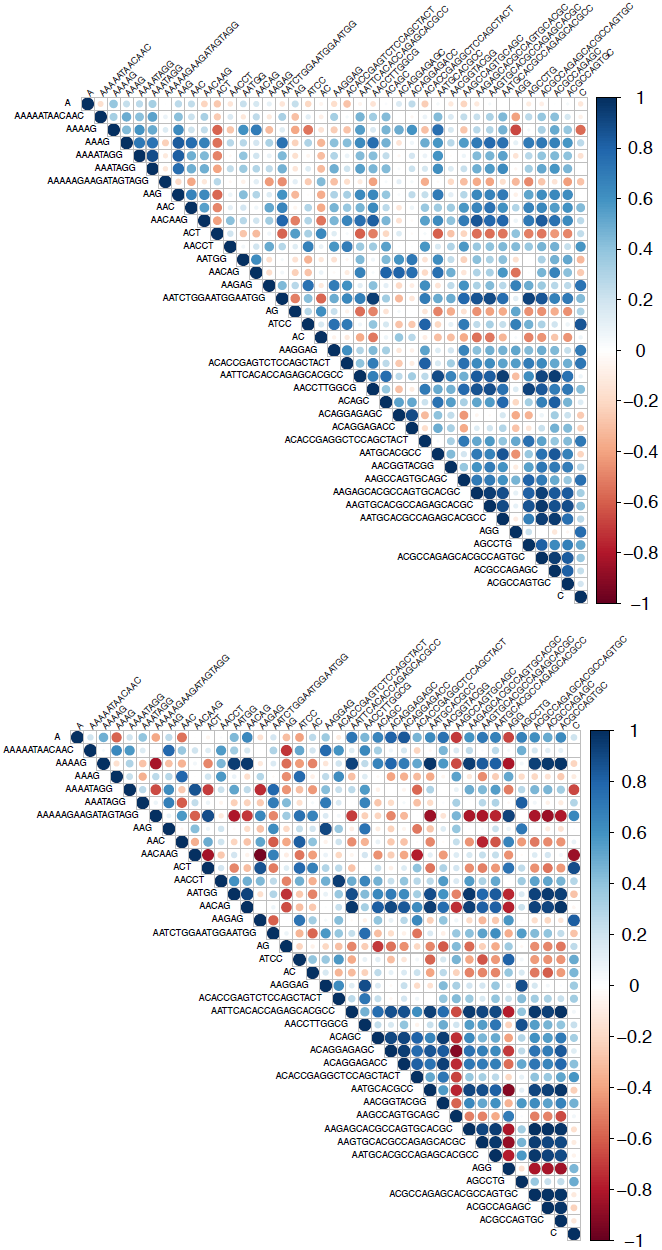
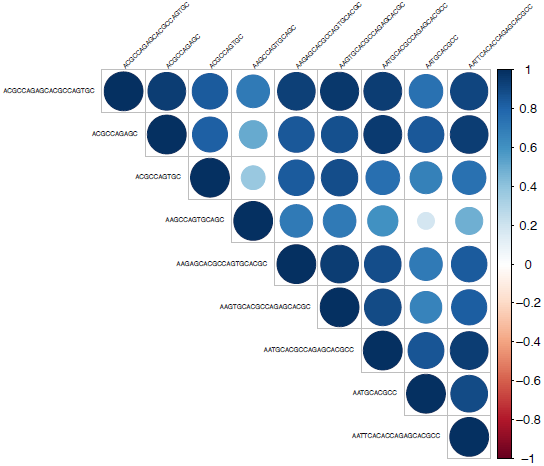
Correlated changes in kmer abundance in (A) the MA lines, and (B) the population. (C) shows the correlations between the family of 9 kmers that are closely related by sequence and GC content, which are situated at the bottom corner of the matrix in B. Matrices were computed from a pairwise Pearson correlation matrix between kmers in the deviation of their abundance from the inferred ancestral abundance. Kmers are ordered by their GC content.

For both the MA lines and the population, strong correlations between kmers that had GC contents between 0.6-0.7 were apparent. Although it is possible that the correlations reflect technical bias, we think this is unlikely because these strong correlations were between the nine kmers closely related in sequence (Figure 2). These nine 10-20 bp kmers have a GC content between 0.6 and 0.7, and their deviations were in fact strongly positively correlated (Figure 5c). Aggregations of kmers that are related in sequence might easily explain some of the patterns of correlation among kmer sequences that we identified. To follow up on this, we analyzed the paired-end reads to measure the level of interspersion between kmer sequences. All kmers were highly interspersed with themselves, indicating the that kmer arrays we are studying span at least the length of the sequenced fragments (~250 bp). Several groups of kmers were interspersed with each other (Supplementary Material, File S5). The telomere repeat, unsurprisingly, was not interspersed with any other of the top 39 kmers. The group of nine related kmers mentioned above were the most mutually interspersed as a group. Eight of the nine were interspersed with at least one other in the group, and the more abundant kmers of the group were interspersed with up to six other kmers within the group.

## DISCUSSION

To our knowledge this is the first study that assays the mutation rates of genome-wide tandemly repeated satellite DNA using MA lines. The inclusion of data from a population that experienced selection also enables us to begin to detect whether selective forces act on satellite DNA. We found that mutation rates in satellite DNA are high, but they appear to be constrained by selection. We assumed the population isolates maintained the ancestral satellite composition when calculating mutation rates, which was reasonable given that the means of the population and MA lines were similar for most satellites, and the population isolates showed narrow changes in kmer abundance. The high variance in the abundance of kmers across the MA lines resulted in high estimated mutation rates. We are confident that these results are biological and not due to technical biases because we normalized kmer counts by the GC-normalized read coverage (see Materials and Methods), and after normalization, there was no longer a correlation between sequencing depth and kmer abundance (Supplementary Material Figure S4). All MA lines and population isolates contained a similar proportion of reads with kmers to total reads (Supplementary Material Figure S1). There was also a high level of concordance in the kmer content between technical replicates. Another line of support that these data are reliable for this type of study is that the analysis of copy number variation of mapped reads revealed mutation rate estimates close to those of previous studies (Chain et al. 2017 in prep). Going forward, using PCR-free library preparation methods can reduce batch-based biases in sequencing results (Wei et al. 2017, in prep).

### High mutation rates can explain the rapid turnover of satellites

The rates of mutation in satellite sequences in our *D. pulex* MA lines were extremely high, ranging from 0.29 to 105 copy changes per generation for a given kmer sequence. Our mutation rates include the sum of changes across all loci of a particular kmer, thus can include both loss/gain of entire repeat arrays (i.e. from recombination), and changes in the number of repeats within individual loci (i.e. from replication slippage). This makes our study distinct from previous studies of microsatellite mutation rates that have focused on single loci (Seyfert et al. 2008). Here, mutation rates estimated on a per-copy basis for satellite repeats we examined were on the order of 10^-3^ copies/generation/copy. Seyfert et al. (2008) estimated the per locus mutation rate of *D. pulex* dinucleotide microsatellites 13-47 repeats long to be on the order of 10^-5^ to 10^-4^ copies/locus/generation. This indicates not only that many satellite sequence arrays are present as multiple loci and long arrays, but that they mutate at much higher rates than microsatellites typically studied. One contributing factor could be that the short microsatellite arrays are more constrained since they are often found in introns or even coding regions (Ellegren 2004). Past studies of microsatellites have found that rates of mutation are positively correlated with the number of repeats, but limited data points prevented the determination of a linear relationship (Wierdl et al. 1997; Brinkmann et al. 1998). We had mutation rate estimates for 39 kmer sequences ranging in abundance from 100 - 50,000 copies and found a significant positive linear correlation between copy number and mutation rate. This suggests a simple relationship between the number of copy units present and the potential for copy units to be gained or lost, consistent with replication slippage being the dominant mechanism of mutation (Schlötterer 2000). We also found that expansions are more common than contractions, indicating that satellite DNA in *Daphnia* has the intrinsic tendency to expand, which is in agreement with the consensus finding in microsatellites across taxa (reviewed in Ellegren 2004).

This high mutation rate demonstrates that there is a high potential for rapid turnover in satellite DNA sequences between species, and even substantial differences in abundances among diverged populations of the same species. Wei et al. (2014) used k-Seek to characterize the satellite sequences in eight different populations of *D. melanogaster*, distributed globally. They found that the overall satellite content varied by a factor of 2.5, accounting for about a 4 Mb difference. This initially surprising finding makes sense in light of our results that the MA lines that were only 82 generations diverged differed in their overall satellite content by a factor of 1.67, accounting for about 0.67 Mb. Even the population isolates, which showed a reduction of variation in satellite content compared to the MA lines, varied in their overall satellite content by a factor of 1.27. This highlights the great potential for expansions and contractions in satellite sequences, which can explain the high levels of differentiation in genome-wide satellite compositions between populations and related species (Wei et al. 2014; Subirana et al. 2015; Wei et al. 2017 in prep).

*Daphnia* and *Drosophila* are both arthropods, although they are estimated to have diverged about 400 million years ago (Rehm et al. 2011). Given the high differentiation observed between closely related species, the paucity of shared sequences and motifs between these taxa is not surprising. It is worth noting, however, that the AGG and AAG motifs as well as a couple AG-rich satellites were conserved between the two species. This conservation could be random or it could indicate a conserved functional role for AG-rich satellites; for example, involvement in binding conserved proteins. The GAGA factor is known to bind to AG-rich satellite sequences in *Drosophila*, although it has been hypothesized to have evolved in the ancestor of Diptera and Hymenoptera and is not thought to be present in *Daphnia* (Heger et al. 2013). However, it is possible that other proteins could be playing a role in binding to AG-rich satellites in *Daphnia* similarly to *Drosophila*. Another possibility is that AG-rich satellites result in a favourable DNA curvature (Palomeque and Lorite 2008). *D. melanogaster* populations were found to be enriched in kmers that were multiples of 5 bp long (i.e. 5, 10, 15, 20, -mers) (Wei et al. 2014). Interestingly, we also found that *Daphnia* were enriched in kmers 5, 10, and 20 bp long. Although there were no 15-mers at high abundance (≥100 copies), 15-mers were enriched when considering the dataset of 162 kmers. This pattern, conserved from *Drosophila* to *Daphnia*, could indicate conservation in the periodicity of DNA wrapping around histones (Lohe and Brutlag 1987).

Based on the new kmers generated *de novo* in the MA experiment and the relatedness of existing kmers, we suggest that novel kmers typically result from a point mutation in an already abundant kmer, followed by expansion of the new kmer. This is in agreement with the result from Wei et al. (2017 in prep), who found new kmers originating in different *Drosophila* species that also differed by only a single nucleotide from a pre-existing kmer. Additionally, satellite kmers can be generated when they expand from a sequence motif that is already present in only a few copies, as we suggest could have occurred for some of our new kmers (Table 2). Concordant with these hypotheses, we found that two kmers were gained independently in two independent MA lines, demonstrating the high potential for new satellite sequences to be generated from existing ones or from an existing motif, and that parallel independent gains of the same satellite are possible. We detected the newly-generated kmers at low abundance, but after many more generations they would presumably have the potential to expand and achieve higher abundance in the genome. Most of the lost kmers were not completely lost, but contracted to be part of a short enough array to not be detected by k-Seek (Table 3). Some of the satellite losses we found likely resulted from deletions of segments of chromosomes. Flynn et al. (2017) found that the MA line C40 experiences homozygous deletions totalling ~100 Mb on chromosome 11. Here we found that C40 lost 3 different satellite sequences, so we think it is likely that these satellites were present on the deleted regions of chromosome 11 (although these tandem sequences were not present in tandem in the genome assembly). In fact, C40 experienced a complicated recombination event that resulted in these deletions and complete loss of heterozygosity across the chromosome (Flynn et al. 2017). It is possible that the satellite DNA composition on this chromosome causes it to be a hotspot for structural rearrangements. Nevertheless, if most of our kmer losses resulted from deletions of large segments also containing functionally important regions, these would likely be purged from natural populations and thus would not be detectable in population data.

### Selection on satellite content

In contrast to the MA lines, which accumulate mutations with minimal regard to their phenotypic consequences, we found that most kmers experienced constraint in their evolution in the non-MA population. The constraint was disproportionately distributed among kmer sequences, with some being unconstrained in these conditions and others being highly constrained. Both the coefficient of variation and mutation rates were lower for many kmers in the population. The population reproduced asexually and it had a census population size of 100-250, indicating selection would have to be substantially strong in order to produce the signal that we observed. Our results suggest that satellite repeats have the intrinsic tendency to expand in the *Daphnia* genome, but the magnitude of expansions are reduced under a selection regime. Although this study does not investigate specific functional roles of satellites, we recognize the potential for some satellites to have specific roles (e.g. Fry and Salser 1977; Palomeque and Lorite 2008; Plohl et al. 2008). This may be true for AG-rich satellites, which are known to bind proteins and are conserved in both *Drosophila* species and *D. pulex* (Raff et al. 1994). The selection we infer might stem from conservation of a specific role for some satellites, and/or deleterious effects of changing the copy number of satellites.

In calculating mutation rates, we assumed that the population maintained the ancestral kmer content. Concordant with this assumption, there was little variation in the total summed abundance of satellite sequences in the population. This would be consistent with the operation of stabilizing selection on overall satellite content (implying that there is an “optimal” level). However, we note that it is possible that negative selection (purging satellite DNA) could have been acting in the population and the lower realized mutation rates in the population was a result of mutation-selection balance (i.e. mutation pressure to increase satellite content, selection to reduce it). Under stabilizing selection, the reduced realized kmer mutation rates in the population could stem from selection on overall genome size or chromosome organization. Previous studies have shown the importance of satellites for maintaining chromatin organization and that genetic variation in satellite content can affect genome stability and chromosome function (Kim et al. 2009, Aldrup-MacDonald et al. 2016). If the genome has evolved to accommodate satellite arrays, altering the length of an array could alter the chromatin organization and disrupt nearby functional regions, causing deleterious effects. This could be especially important for *Daphnia*, which has a compact genome with little intergenic space (Colbourne et al. 2011). If selection to remove satellite DNA was the dominant form of selection, this would imply that the satellite arrays were mostly “junk” and the genomes would be in a constant state of purging satellite DNA. Moreover, it seems that the level of selection varies across kmers. We found five kmers to be under very little constraint, shown by the lack of difference between their CV and mutation rates between the MA lines and population (Figure 4 c and d).

### Correlations in kmer changes

Across the MA lines, we found mainly positive correlations between the kmers in their expansions and contractions. These could have arisen randomly or been driven by physical location/linkage. We found that kmers that were closely related to each other were strongly correlated in their changes in abundance, which was especially evident for the family of 10-20 bp kmers with the GCCAG motif (Figure 2, top left). Concordant with the hypothesis that new satellite sequences come from point mutations in existing ones, we found that these closely related kmers were highly interspersed within each other. Since these positive correlations were present in both the MA lines and the population, it is likely to be a consequence of the mutational process. Segmental duplications and deletions, which are common in *Daphnia* (Keith et al. 2016), could explain the observed positive correlations. When only the population isolates were considered in the correlations, we identified more correlations of stronger effect, both positive and negative. The positive correlations could arise from the same selective regime acting on pairs of kmers, i.e. if expansions of both kmers of the pair in question are both selected against. On the other hand, the negative correlations may arise if different satellite sequences are in conflict (Wei et al. 2014). Specifically, there could be selection on a chromosome to maintain its organization such that if one kmer happens to expand, another will have to contract. Kmers localized on the same chromosomal region may be the ones negatively correlated with each other, however, we were not able to test this thoroughly with our data because of the un-mappable nature of repeats.

## Conclusions

We were able to gain a picture of genome-wide satellite DNA expansions and contractions, and provide evidence that satellite DNA is not always neutrally evolving but can experience strong selection. We also show evidence that new satellite sequences are often generated from existing ones, and there is a complex interaction structure between individual satellites that differs if selection is at play. Future studies using different sequencing technologies, especially with longer reads that can capture longer satellite sequences and span the boundaries of unique and repeated sequences, will provide further insights into satellite DNA evolution.

## ACKNOWLEDGEMENTS

We would like to thank D. Barbash and F. Chain for useful discussions, and K. Wei for providing scripts for the kmer interspersion analysis. J. Bull provided summary data from the life history experiment that allowed us to calculate the number generations in the non-MA population. We would also like to thank S. Lower for helpful discussions on the analysis of the data and comments that helped improve the quality of the manuscript. The original MA lines and sequencing was supported by NSERC Discovery Grant (341423-2012) to MEC and the kmer analysis was supported by NIH grant R01 GM119125 to AGC and D. Barbash. JMF was supported by a NSERC PGS-D scholarship.

